# Environmental responsiveness of flowering time in cassava genotypes and associated transcriptome changes

**DOI:** 10.1101/2021.02.11.430817

**Authors:** Deborah N. Oluwasanya, Andreas Gisel, Livia Stavolone, Tim L. Setter

## Abstract

Cassava is an important food security crop in tropical regions of the world. Cassava improvement by breeding is limited by its delayed and poor production of flowers, such that cassava flowering under field conditions indirectly lengthens the breeding cycle. By studying genotype and environment interaction under two Nigerian field conditions (Ubiaja and Ibadan) and three controlled temperature conditions (22°C/18°C, 28/24°C and 34/30°C (day/night)), we found that while early flowering genotypes flowered at similar times and rates under all growing conditions (unfavorable and favorable field and controlled-temperature environments), late flowering genotypes were environmentally sensitive such that they were substantially delayed in unfavorable environments. Flowering times of late genotypes approached the flowering time of early flowering genotypes under relatively cool Ubiaja field conditions and in growth chambers at 22°C, whereas warmer temperatures elicited a delaying effect. Analysis of field and controlled temperature transcriptomes in leaves revealed that conditions that promote early flowering in cassava have low expression of the flowering repressor gene TEMPRANILLO 1 (TEM1), before and after flowering, among others. Field transcriptomes showed that the balance between flower promoting and inhibitory signaling, appeared to correlate with flowering time across the environments and genotypes.

## 1 Introduction

Cassava is a tropical plant originating from the Amazonian region, which is cultivated for its starchy storage roots [1]. It is an important staple food in the tropics and ranks as the fifth most important source of starch in the world [2]. Although it can be propagated asexually, to develop improved cultivars through breeding requires sexual reproduction and associated genetic recombination and selection for genetically superior traits [3]

Sexual reproduction in cassava is limited at multiple phenological stages ranging from the transition to flowering to the development of fruits and seeds [4–8]. Flowering time is a very critical factor in cassava’s sexual reproduction because it determines the length of the cassava breeding cycle. The development of new cultivars takes eight to ten years [9] due to the difficulty of genetic recombination caused by delayed flowering or no flowering at all [3]. Cassava flower development is associated with the development of fork type branches (sympodial branching) and in this paper forking is synonymous to flowering. A more detailed description of cassava’s reproductive development has been provided by [7]

Flowering induction is regulated by environmental cues (such as temperature and photoperiod) to ensure that flowering occurs under the most optimal conditions for reproductive success [10]. The role of temperature in regulating flowering time is particularly important for cassava that is grown in the tropics where daylengths do not vary significantly throughout the year. In cassava, flowering time is favored by long days and relatively cool (but not vernalization) temperatures [5]. This is in contrast to the model plant *Arabidopsis thaliana*, in which the time to flowering is hastened in warmer ambient temperatures [11], although it also flowers in response to long days (short nights) [12]. The genetic control of flowering time has been well characterized in *A. thaliana* and over 300 genes have been identified by forward and reverse genetics to be involved in flowering time regulation, as documented in the flowering database FLOR-ID [13]. The Flowering Locus T (FT) gene has been shown to be a flowering integrator of multiple flower inductive pathways and is positively correlated with flowering time in most species studied so far [14]. In cassava, as in many other species, the overexpression of the Arabidopsis FT gene and native cassava FT gene has been shown to accelerate flowering time in otherwise very late flowering genotypes [4, 15, 16]. This provides strong evidence for the involvement of FT in regulating also cassava flowering time. In Arabidopsis, warm temperatures are favorable for flower induction and, correspondingly, FT expression is elicited by long days and warmer temperatures [11]. In cassava plants expressing two homologs of FT, MeFT1 and MeFT2 [5], flowering is stimulated by long days, and correspondingly, long days elicit expression of MeFT2 at the end of long days. However, while cool temperatures are favorable to flower induction, increased expression of MeFT1 and MeFT2 in response to cool temperature is not consistent among genotypes, suggesting that other signaling factors, such as inhibitory factors, might be involved [5].

Researchers at the International Institute of Tropical Agriculture (IITA), Ibadan, Nigeria had previously identified a field location (Ubiaja, Nigeria) under which flowering occurs earlier and general flower development is enhanced [17, 18], independently of the soil characteristics of the two environments [18]. Consistently with earlier cassava flowering, weather reports show that temperature is generally cooler in Ubiaja.

We compared the flowering behaviors of eight genotypes (representing a range of flowering times) under the ‘favourable’ Ubiaja field and the ‘unfavorable’ Ibadan field conditions. In parallel, we studied the effect of temperatures (ranging from 22°C to 34°C) on the flowering times of a subset of these genotypes comprising three genotypes under controlled conditions. Finally, the transcriptome analysis of a selected pair of genotypes before forking and seven days after forking, both in the field and in growth chamber conditions, demonstrated that the expression of a group of flowering-related genes is consistently regulated under favorable and unfavorable flowering conditions.

In the face of the world climate change challenges, understanding the molecular basis of flowering time control in cassava is critical to enhance cassava breeding for crop improvement and opens new possibilities to develop strategies and methodologies to allow cassava flowering irrespective of the environmental growth conditions.

## 2 Materials and methods

### 2.1 Plant Materials and Growing Conditions

#### (a) Field Station

Field experiments were conducted from June 2017 to January 2018 at two field stations in Nigeria: Ibadan (7.4° N and 3.9°E, 230 m asl) in Oyo State and Ubiaja (6.6° N and 6.4° E, 221 m asl) in Edo State. Cassava stems of similar lengths (about 20 cm each), were planted simultaneously in June 2017 at both locations so ages of plants were identical. Eight genotypes were selected from the IITA diversity population named the Genetic Gain Population. These genotypes were selected based on previous information about their flowering times. Three categories were selected for our study, namely (i) early flowering (< 60 days after planting [DAP]), represented by IITA-TMS-IBA010615 and IITA-TMS-IBA020516, (ii) middle (60 – 99 DAP), represented by IITA-TMS-IBA030275, IITA-TMS-IBA010085, and IITA-TMS-IBA980002, and (iii) late (> 100 DAP), represented by IITA-TMS-IBA8902195, IITA-TMS-IBA000350, and TMEB419. They are available from the IITA germplasm bank (Ibadan, Nigeria; accession list: https://www.cassavabase.org/accession_usage). In this manuscript, these genotypes will be referred to as ‘615, ‘516, ‘275, ‘085, ‘0002, ‘2195, ‘350, and ‘419, respectively.

Plants in each location were grown in a randomized block design consisting of 6 blocks each with the eight genotypes randomly assigned as plots. Each plot contained 8 plants grown in a 2×4 matrix at 1m x 1m spacing.

#### (b) Growth Chamber

One early genotype - IITA-TMS-IBA020516 and two late genotypes - IITA-TMS-IBA8902195 and IITA-TMS-IBA000350, were grown in tissue culture at the Genetic Resources Center, International Institute of Tropical Agriculture, Nigeria. Plantlets were screened to ensure absence of infection and other appropriate phytosanitary conditions. Tissue culture plants were shipped to Cornell University, Ithaca, NY, USA and were transplanted to soil and grown several months to form plants with stems >15 mm diameter. Stakes of about 15 cm length were cut from the stems of established plants and used as propagules for experiments. Plants were grown in three growth chambers set at 22°C/18°C, 28°C/24°C, and 34°C/30°C, day/ night temperatures, respectively. Photoperiod was held constant at 12 h light and 12 h dark. Plants were completely randomized in each growth chamber. Each chamber had two replicates of each genotype. Two independent batches of this experiment were carried out. Growth chambers were Conviron Controlled Environments, Ltd (Winnipeg, Manitoba, Canada) model PGW 36 walk-in growth rooms (135 × 245 × 180 cm [ht.]) with ten 400 W high pressure sodium and ten 400 W metal halide lamps providing 600 μmol photons (400-700 nm) m^-2^ s^-1^. Root-zone potting mix and fertilization were as previously described [5].

### 2.2 Data collection

At Ubiaja and Ibadan, daily temperature and rainfall were collected by temperature loggers, Onset ^®^ HOBO Pendant (https://www.onsetcomp.com/products/data-loggers/mx2202, Bourne, MA, USA) placed in ventilated reflective shelters [19] at 1.1 m height and by an automated self-emptying rain gauge – RainWise ^®^ (https://rainwise.com/rainlogger-complete-system, <u>Trenton, ME, USA</u>). Flowering time was recorded as the time (DAP) of appearance of the first reproductive branching (forking). Number of nodes was counted from the soil surface to first fork on each plant. Plant height, whole plant fresh weight, storage root fresh weight and number of storage roots were recorded at 7 months after planting on the field and growth chamber. Data was collected using Field Book software application [20]

### 2.3 Statistical Analyses

Field Data was modelled using a linear mixed model while growth chamber data was modelled using a simple linear model. In the field study, locations and genotypes were fixed effects, while blocks were random effects. In the growth chamber study, temperature (T), genotype (G), and T *×* G interaction were the modelled sources of variation. Both models were tested by analysis of variance. Flowering time and fraction of plants flowered were subjected to survival analysis using the Kaplan-Meier’s curve [21]. Multiple means comparison was conducted in the emmeans package [22] using the Tukey-HSD method. All analyses were conducted in R [23].

### 2.4 Transcriptomic Analysis

Genotypes ‘0002 and ‘419 were selected for field transcriptomics while genotypes ‘516, ‘350, ‘2195 were selected for controlled temperature transcriptomics. These genotypes represented the range of early and late flowering lines with varying degrees of environmental responsiveness. Leaf tissue samples were collected from the youngest fully expanded leaf on each plant. Three and five biological replicates were collected from field and growth chamber plants, respectively. The field samples were collected at 21 DAP (preforking) and 7d post forking (relative to genotype development). In the growth chamber, samples were collected at 47 and 96 DAP. Samples were obtained in the late afternoon (Ubiaja and Ibadan) or within 1.5 h of the end-of-light period (growth chambers) and immediately placed in porous polyester tea bags and immersed in liquid N_2_ to freeze and for storage.

Total RNA was extracted from each sample by a modified CTAB protocol. For field samples about 0.2g of frozen leaf tissue were ground with mortar and pestle after which it was transferred to 1.5ml Eppendorf tubes to which 1 mL of preheated (65°C) CTAB extraction buffer was added (Buffer comprised of 2% [w/v] CTAB detergent, autoclaved 0.1M Tris-HCl pH 8, 20mM EDTA, 1.4M NaCl and 2% PVP, with pH adjusted to 8.0). Samples were warmed at 65°C for 15 mins with vortexing at 5-minute interval after which they were centrifuged at maximum speed for 5 minutes. To 1 mL of supernatant in a fresh Eppendorf tube, 1 mL of chloroform Isoamyl alcohol (24:1) was added, vortexed and centrifuged for 10 min. Supernatant was collected in a clean Eppendorf tube to which cold 2-propanol was added (0.6 volume of supernatant) and mixed by inverting gently. Samples were centrifuged for 10 min at maximum speed to collect pellets which were washed in 70% ethanol and air dried. Pellets were redissolved in RNase free water, treated with DNase I and cleaned with RNA Clean and Concentrator (Zymogen). RNA quality was determined by gel electrophoresis and RNA was bound to matrix in RNAstable^®^ and shipped to Cornell University, Ithaca, NY. RNase free water was added to RNAstable^®^ to recover RNA for downstream assay. Growth chamber samples were ground to a fine powder in a mortar and pestle chilled with liquid N_2_; about 0.5 g of the powder was vigorously mixed for 5 min with 1 mL of CTAB extraction buffer; 0.2 mL of chloroform was added and mixed for 15 s, tubes were centrifuged at 14,000 g for 10 min and the top layer was removed to a new tube. To these samples was added 700 μL of Guanidine Buffer (4M guanidine thiocyanate, 10 mM MOPS, pH 6.7) and 500 μL of ethanol (100%). This mixture was applied to a silica RNA column (RNA mini spin column, Epoch Life Science, Missouri City, TX, USA), then alternately centrifuged and washed with 750 μL of 1) Tris-ethanol buffer (10 mM Tris-HCl [pH 7.6], 1 mM EDTA, containing 80% [v/v] ethanol), 2) 80% ethanol (twice), and 3) 15 μL RNAase-free water (to elute the RNA). The RNA quality of field and growth chamber samples were evaluated for quality with a gel system (TapeStation 2200, Agilent Technologies, Santa Clara, CA, USA). Other downstream assays were same for both field and growth chamber samples.

cDNA libraries were prepared using the Lexogen Quantseq FWD kit [24] and DNA was sequenced by the 3’ RNASeq method [25] using an Illumina NextSeq500 sequencer at the Genomics Facility, Cornell Institute for Biotechnology. Software was used to remove Illumina adapters, poly-A tails, poly-G stretches [26]. The trimmed reads were aligned to the Manihot esculenta genome assembly 520_v7 using the STAR aligner (version 2.7.0f) [27].

Differential Gene expression analysis was conducted using the DESeq2 package by Bioconductor [28]. Each transcript was annotated by the best match between Manihot esculenta genome v7 and the Arabidopsis genome as presented at Phytozome13 [29].

Gene ontology and enrichment analysis were carried out using the ShinyGO app (http://bioinformatics.sdstate.edu/go/) [30]. A combined list of Arabidopsis flowering genes were obtained from the Max Planck Institute (https://www.mpipz.mpg.de/14637/Arabidopsis_flowering_genes) and Flowering Interactive Database (FLOR-ID) (http://www.phytosystems.ulg.ac.be/florid/) [31] and a list of hormone signaling genes sourced through the Database for Annotation, Visualization and Integrated Discovery (DAVID) (https://david.ncifcrf.gov/) [32] were used to examine the expression profiles of flowering and hormone signaling genes.

## 3 Results

### 3.1 Field Experiment

#### 3.1.1 Weather

Weather data collected from field sites in Ibadan and Ubiaja are shown in Figure 1. Cumulative rainfall at the two sites were similar in the first month, then diverged for the next two months with Ubiaja receiving more rainfall than Ibadan (Figure1a). Day-time temperatures, as indicated by daily maxima, were generally cooler in Ubiaja than Ibadan with the largest temperature difference in the shaded and ventilated shelters housing the weather instrumentation did not exceed 3 °C (Figure 1b). Nighttime temperatures were essentially the same at the two sites.

**Figure 1.**
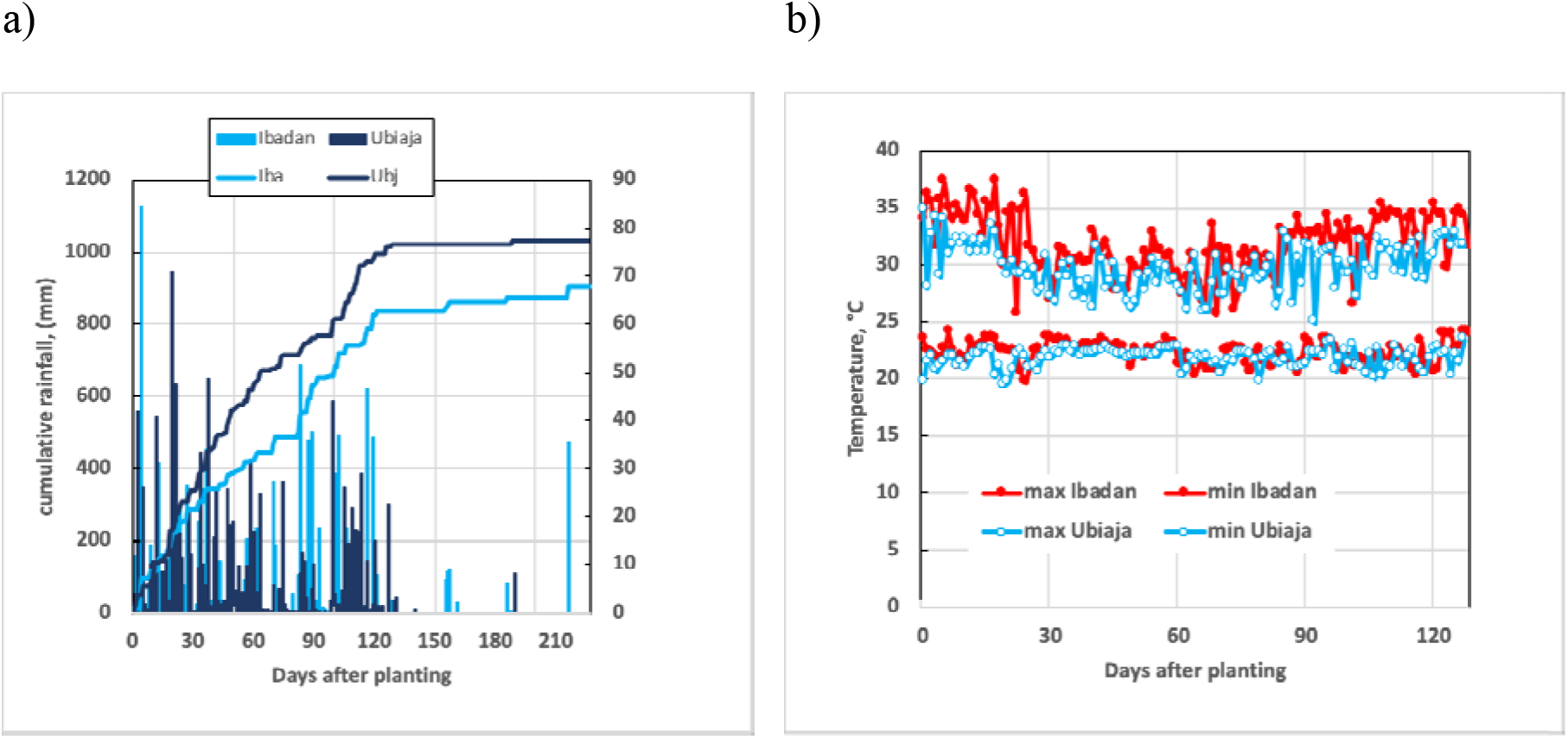
Rainfall (mm) and temperature (°C) collected in 2017 for Ubiaja and Ibadan a) Cumulative rainfall and by events b) Maximum and Minimum daily temperatures.

#### 3.1.2 Vegetative growth patterns under field environments of Ibadan and Ubiaja

Survival of plants generally differed between field environments (Figure 2a). For all eight genotypes, plants grown in Ibadan were between 80 and 130% taller than plants grown in Ubiaja (Figure 2b). The partitioning index (i.e. storage root weight/total plant weight on a fresh weight basis) was significantly higher in Ubiaja than Ibadan, differing by at least 15% between locations (Figure 2c). The lower partitioning index in Ibadan was due to substantially greater above ground fresh weight (about double) than in Ubiaja (Figure S1a). Storage root fresh weight tended to also be greater in Ibadan with some genotypes having as much as 90% higher storage root fresh weight (Figure S1b). Similarly, storage root numbers were significantly higher for all genotypes in Ibadan relative to Ubiaja (Figure 2d). On average across all genotypes, there was 1.7-fold more storage roots in Ibadan than Ubiaja. This increase was similar to the 2-fold higher above-ground fresh weight in Ibadan relative to Ubiaja (Figure 2a). The pattern of vegetative growth between both field locations shows that plants were generally larger and more vigorous in Ibadan.

**Figure 2.**
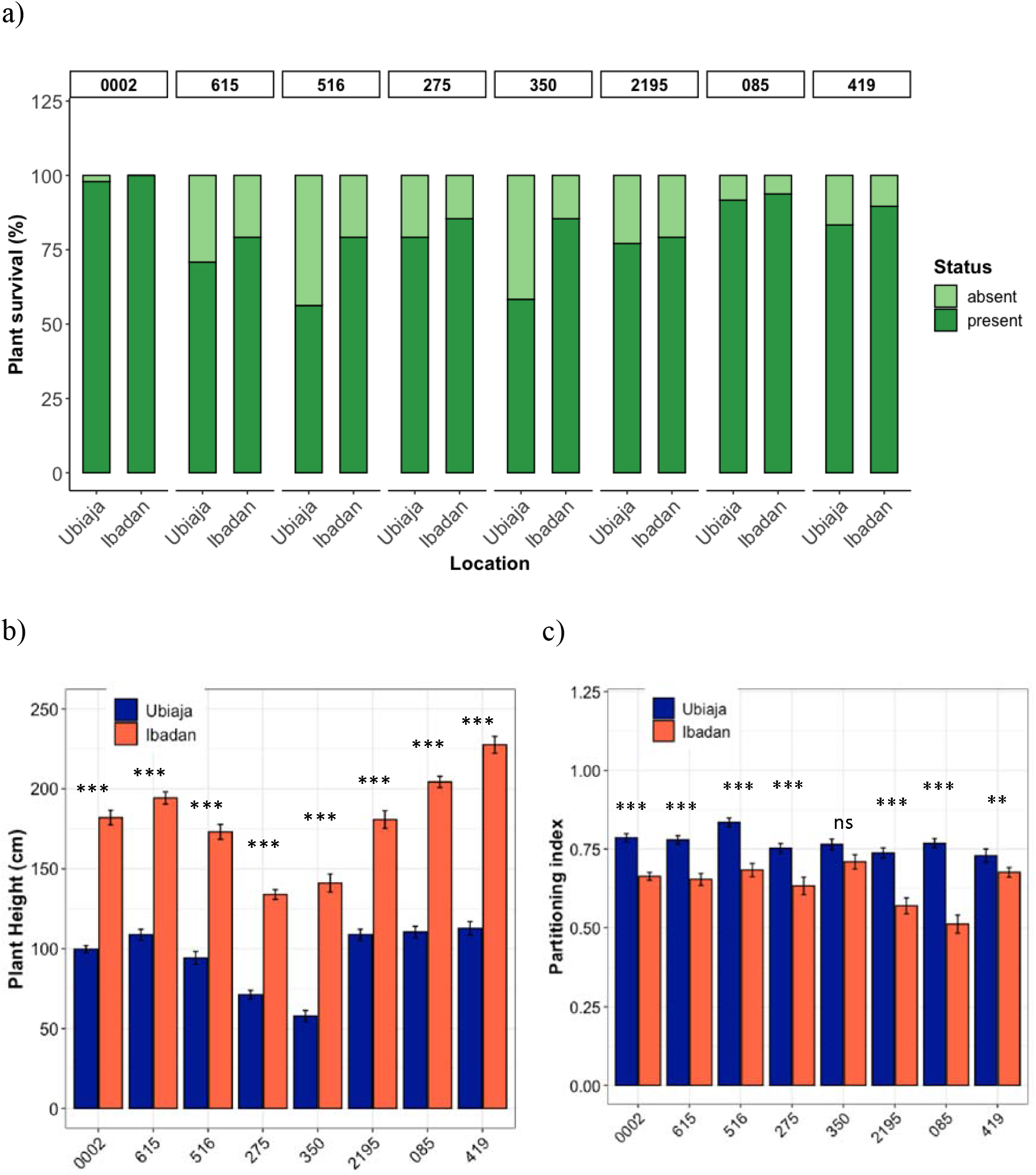
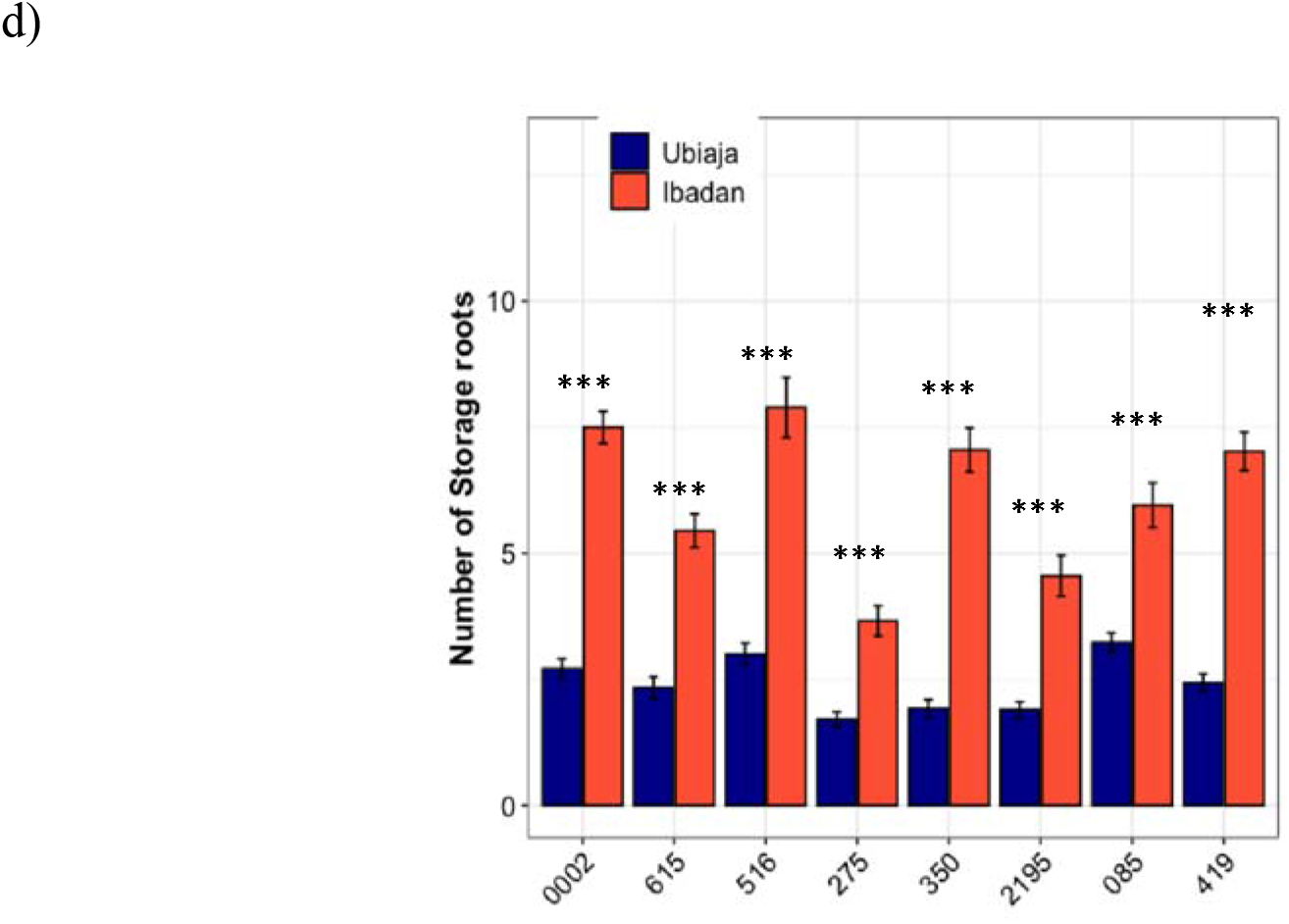
Vegetative growth under field conditions a) Percent plant survival in Ubiaja and Ibadan b) Plant Height (cm) measured from soil surface to highest shoot apex. c) Partitioning index (storage root FW/total plant FW). d) Number of storage roots. *, ** and *** indicate statistical significance on pairwise comparisons between locations for each genotype at 0.05, 0.01, and 0.001 significance levels. Mean partitioning index are reported while data were third order transformed (cubes) for statistical analysis.

#### 3.1.3 Flowering phenotype under field environments of Ibadan and Ubiaja

Using the flowering time in plants that attained flowering within experimental period (i.e., observed flowering time) or age of plants surviving to the end of experiment that did not flower, we plotted Kaplan Meier curves showing the probability of flowering (or the decline in the probability of not flowering) as a function of time (Figure 3a). Genotypes differed in their flowering response between the two locations. Genotypes ‘2195, ‘085, and ‘419 tended to be earlier in Ubiaja. For some genotypes (‘0002, ‘615, and ‘275), rates of progress in flowering with time were nearly identical between locations. In Ibadan, in most of the lines almost all the plants eventually flowered; however, in Ubiaja for ‘0002, ‘275, ‘350 and ‘419 between 10 and 15% of the lines failed to flower during the period of observation (up to 200 dap) (Figure 3a). This phenomenon resulted in cross overs of the flowering curves late in the season (Figure 3a). Genotypes ‘516 and ‘350 were unique because flowering was slightly (but not significantly) delayed in Ubiaja by chronological age (DAP). To provide another measure of the developmental time until flowering, we counted the number of nodes from the soil surface to the flowering fork. There were large differences in the number of nodes for genotypes ‘350, ‘2195, ‘085, and ‘419 and smaller differences in genotypes ‘0002, ‘615, and ‘275 (Figure 3b). In both cases plants in Ibadan had more nodes, indicating later flowering. The number of nodes to flowering were not significantly different between locations in genotype ‘516 (Figure 3c). In Ubiaja, the number of nodes to flowering were relatively similar in all lines (early and late lines averaged 16 and 19 nodes, respectively). In contrast, genotypes differed substantially in Ibadan, (early and late lines flowered averaged 18 and 35 nodes, respectively). It is noteworthy that these flowering differences in genotypic response to environment were not correlated with the differences in shoot or root growth, or partitioning index in Ubiaja and Ibadan (Figure 2c), showing that across this set of genotypes, flowering was not closely related to resource availability, vegetative growth, or partitioning.

**Figure 3.**
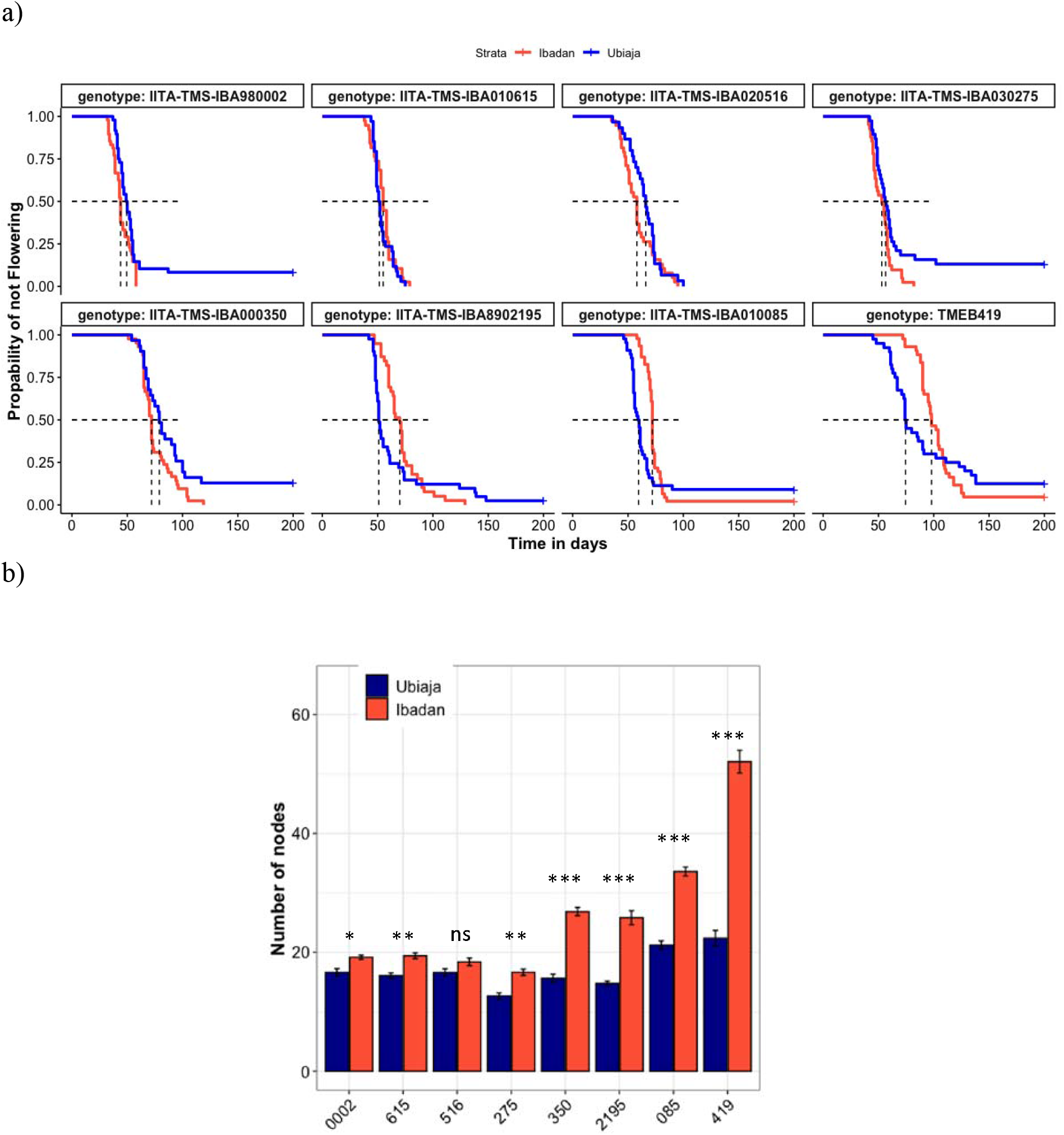
Flowering time responses in Ubiaja and Ibadan a) Kaplan-Meier curves of distinct genotype flowering times in field locations. b) Number of nodes on main stem (counted from soil surface to last node before fork branch) as a developmental time score on field. *, ** and *** indicate statistical significance on pairwise comparisons between locations for each genotype at 0.05, 0.01, and 0.001 significance levels. Mean number of nodes are reported while data was square root transformed for statistical analysis.

### 3.2 Controlled temperature experiment

Given that one of the environmental differences between the field locations was temperature, especially day-time temperature, we tested a set of genotypes for their flowering response to three temperatures in growth chambers. The genotypes selected for this experiment represented the range of response: ‘350 and ‘2195, which developed many more nodes in Ibadan than Ubiaja before flowering, and ‘516, which flowered after approximately the same number of nodes in both environments. In the growth chamber, there was a 100% survival for genotypes.

#### 3.2.1 Vegetative growth patterns under controlled temperatures

In all three genotypes, plant height increased substantially with temperature from 22°C to 34°C (Figure 4a). There were no differences in the partitioning index at all temperatures for genotype ‘350 (similar to field observation (Figure 2c)), none for genotypes ‘516 and ‘2195 between 22°C and 28°C, but a significant reduction between 28°C and 34°C (Figure 4b). In addition, the partitioning index was significantly reduced in genotype ‘2195 between 22°C and 34°C. Overall, the partitioning index tended to be highest at 28°C. This observation in partitioning index arose from the above ground fresh weight being less responsive to temperatures (Figure S2a); while storage root fresh weight increased between 22°C and 28°C; and either plateaued or declined between 28°C and 34°C (Figure S2b). Warmer temperatures in the growth chamber tended to decrease the number of storage roots per plant, especially between 22°C and 34°C where effect was significant (P≤0.05) for ‘516 and ‘350 (Figure 4c).

**Figure 4.**
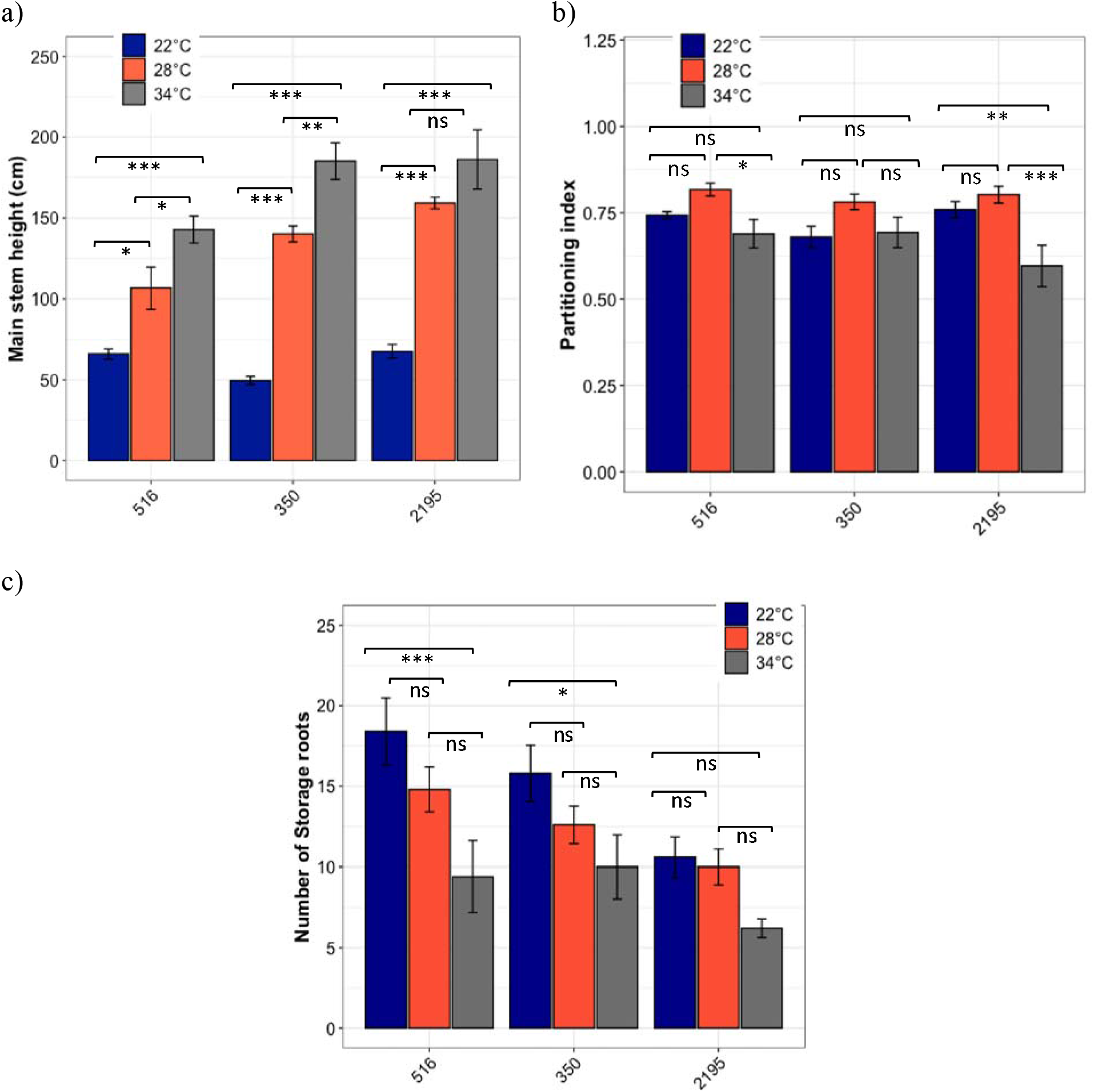
Vegetative growth under controlled temperatures. a) Main stem height (cm) measured from soil surface to highest point in plant before forking. b) Partitioning index (storage root FW/total plant FW) c) Number of storage roots *, ** and *** indicate statistical significance on pairwise comparisons between temperatures for each genotype at 0.05, 0.01, and 0.001 significance levels.

#### 3.2.2 Flowering phenotype under controlled temperatures

In the growth chamber we modelled the probability of flowering (or the decline in the probability of not flowering) as a function of time using the Kaplan Meier method, from the observed flowering time (where flowering occurred within the duration of experiment) or the maximum duration of experiment for plants that did not attain flowering (Figure 5a). At 22°C, all genotypes attained 100% of the plants flowering. In contrast, at warmer temperatures (28 and 34°C) only genotype ‘516 attained 100% flowering, and its flowering was only slightly delayed at 28°C (Figure 5a). In contrast to the other two genotypes, ‘516 had little response to the delaying effect of warmer temperatures. Genotypes ‘350 and ‘2195, however, flowered poorly at warmer temperatures – flowering was completely absent at 34°C, while flowering was between 20 and 30% at 28°C within the period of experiment (Figure 5a). Data on the number of nodes to forking, an alternative measure of developmental timing, confirmed the genotypic differences in temperature responsiveness. The number of nodes to forking in ‘516 did not differ statistically (P≤0.05) amongst temperatures (Figure 5b), confirming this genotype’s insensitivity to a delaying effect by warm temperatures. This finding was analogous to ‘516’s insensitivity of number of nodes to fork among the different environments of Ibadan and Ubiaja (Figure 3b). In contrast, using nodes to fork (or maximum number of nodes countable) as an index of development, flowering in ‘350 and ‘2195 was substantially delayed at warmer temperatures. These genotypes flowered after significantly (P≤0.01) fewer nodes at 22°C than at 28°C and 34°C where flowering was partial or completely absent (Figure 5b).

**Figure 5.**
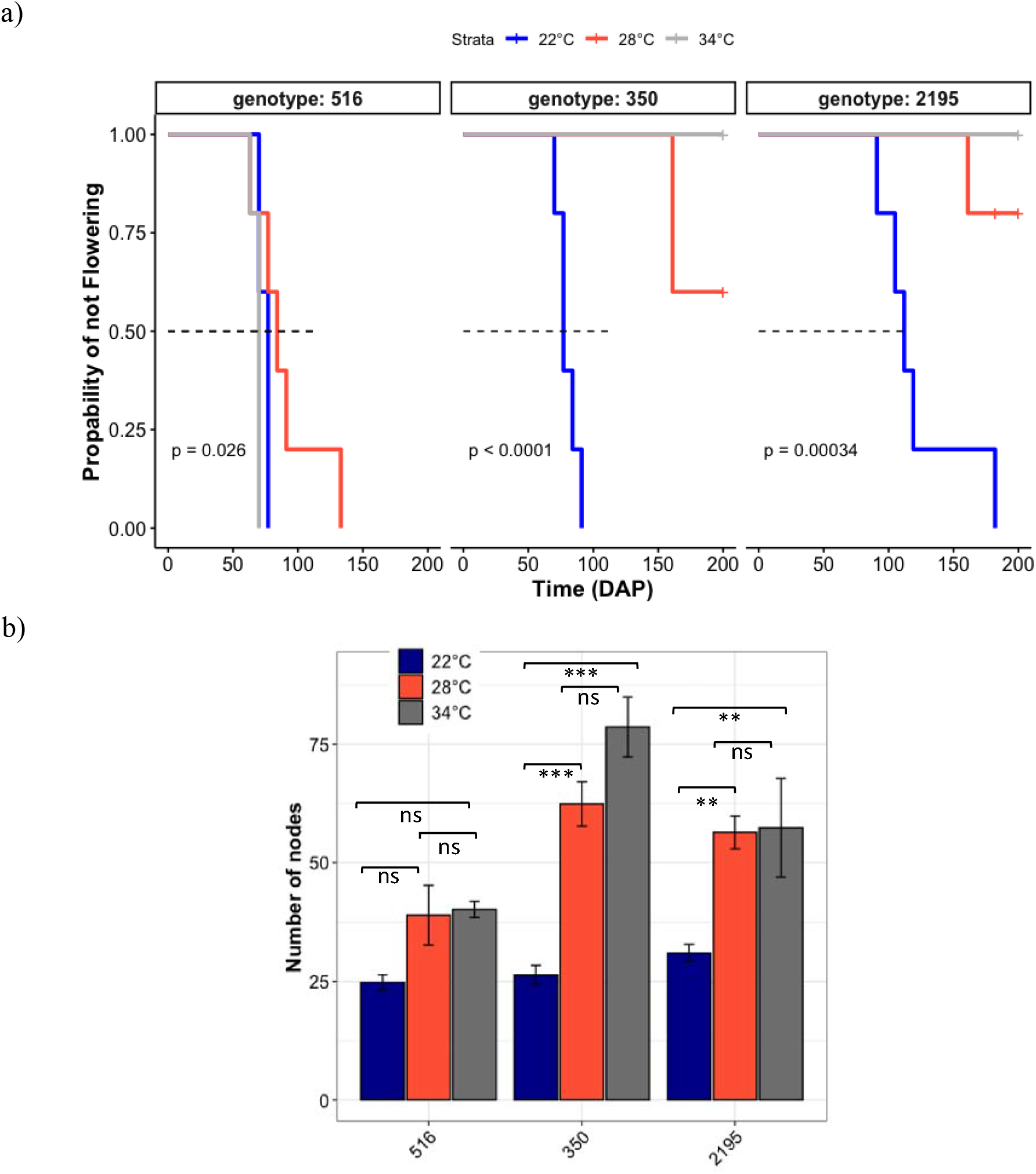
Flowering time responses to controlled temperatures a) Kaplan-Meier curves of distinct genotype flowering times at different temperatures. b) Number of nodes on main stem (counted from soil surface to last node before fork branch or maximum countable where no forking occurred). *, ** and *** indicate statistical significance on pairwise comparisons between locations for each genotype at 0.05, 0.01, and 0.001 significance levels.

### 3.3 Transcriptomics

Transcriptomes were analyzed in mature leaves of plants from our two studies (Field Environments and Growth Chamber) with respect to the following variables: 1) <u>Environment</u> (Ubiaja versus Ibadan in the field study; three temperatures in the growth chamber study); 2) <u>Stage of plant development relative to flowering</u>, where the early stage was before flowering and the later stage was post flower appearance, and 3) <u>Genotype</u>, where lines were chosen to represent a range of environmental responsiveness and earliness of flowering. Samples with fewer than 150,000 demultiplexed reads (were excluded from the gene counting analysis. For the remaining samples, Illumina adapters, Poly-A tails and poly-G stretches were removed. Reads with at least 18 bases in length after trimming were kept.

#### 3.3.1 Field Transcriptome

Under field conditions, in mature leaves for the combined genotypes and sampling dates, 1074 genes were differentially expressed between the two locations with Ibadan as reference. At 5% FDR, 390 genes had higher expression in Ubiaja while 684 genes had lower expression in Ubiaja (Figure 6). Enrichment analysis indicated that the categories of genes that were significantly overrepresented among the genes that had higher expression in Ibadan than Ubiaja were several that relate to abiotic environmental stress, including “Response to abiotic stimulus” (85 genes, p=8.97e-7), “Response to abscisic acid” (29 genes, p=1.81e-3), and “Response to ethylene” (25 genes, 5.74e-7).

**Figure 6.**
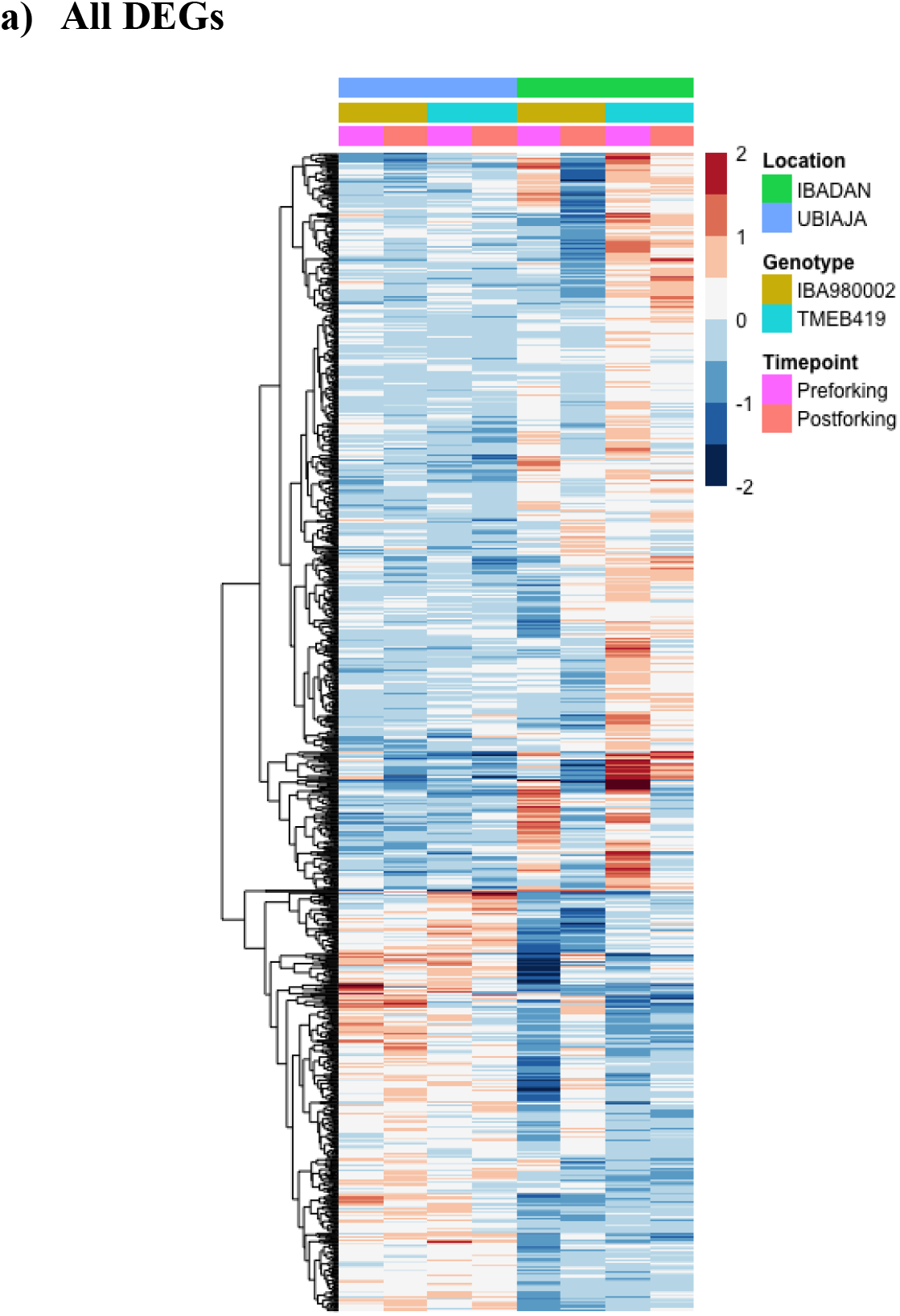
Heat map showing relative expression of differentially expressed genes in response to field location (Ibadan vs Ubiaja), genotype (‘0002 vs ‘419), and timepoint of development (preforking and postforking). Figure shows 1074 genes significantly differentially expressed (α = 0.05) (averages of biological replicates per time point, genotype and location). Colour scale indicates log2 Fold Changes.

##### Flowering time genes

Although the leaf transcriptome in this study is likely to have numerous differentially expressed genes among the tested environments for factors that relate to leaf stress, photosynthesis and metabolic processes, we focused our analysis on genes related to flowering and related signaling. From a list of 240 flowering time genes (see Materials and Methods), nine flowering time genes were differentially expressed in the field transcriptome (Figure 7a). These genes generally showed location sensitivity and had similar expression profiles at the postforking stage (Figure 7a).

**Figure 7.**
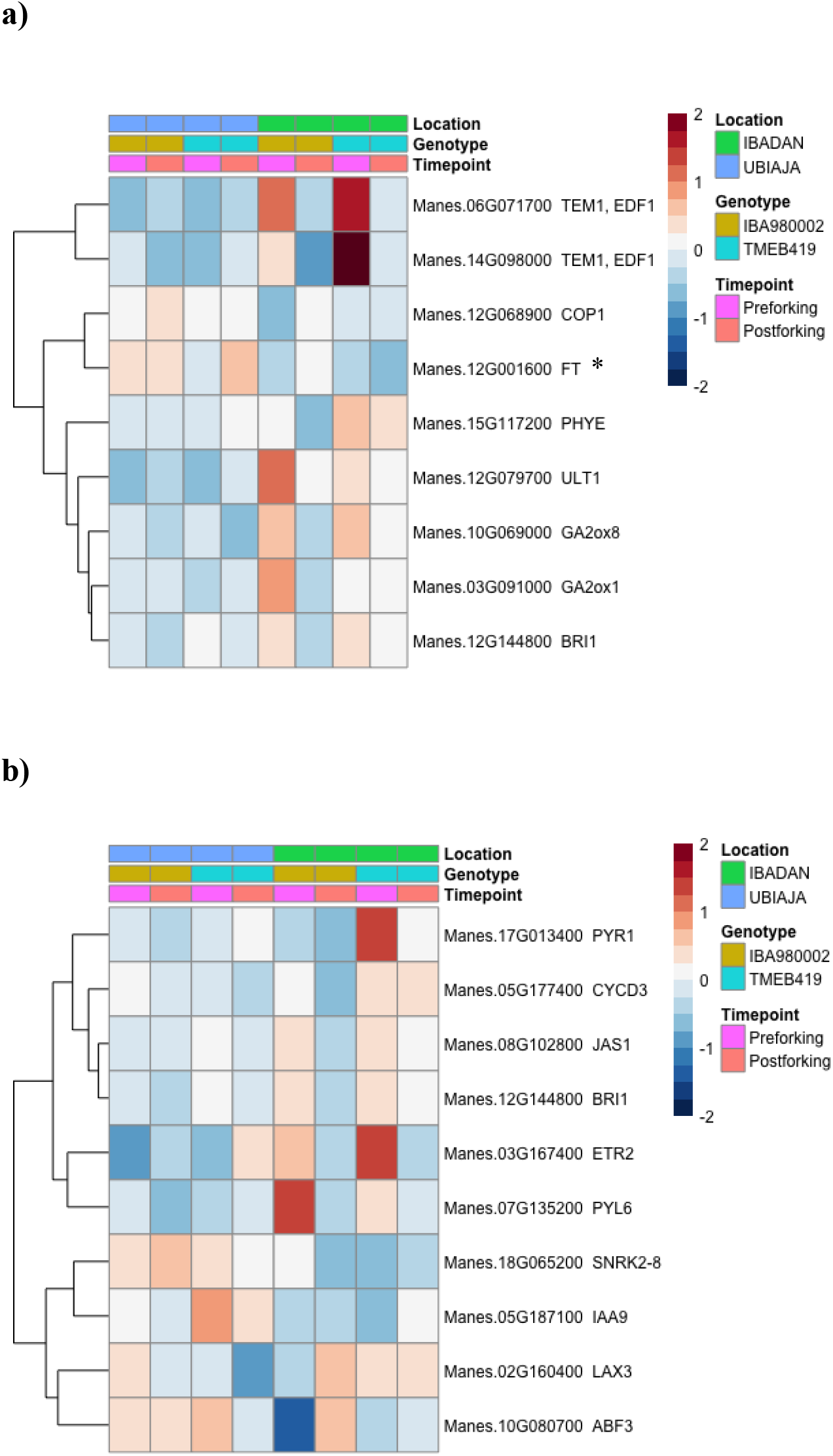
Flowering time and hormone signaling genes differentially expressed under field conditions. The heat map shows relative expression across location (Ubiaja vs Ibadan), genotype (‘0002 vs ‘419), and timepoint of development (preforking and postforking) a) Flowering time genes b) Hormone signaling genes *Cassava homologue of Arabidopsis FT gene – Manes.12G001600 has been named MeFT1 as in (Adeyemo et al. 2018; Odipio et al. 2020)

##### Hormone signaling genes

From a list of 160 select hormone signaling genes spread across eight plant hormones (see Materials and Methods), 10 hormone signaling genes were differentially expressed in the field transcriptome (Figure 7b). These genes were involved in abscisic acid (4 genes: PYR1, PYL6, SNRK2-8, ABF3), auxin (2 genes: IAA9, LAX3), cytokinin (CYCD3), ethylene (ETR2), jasmonic acid (JAS1) and brassinosteroid signaling (BRI1).

#### 3.3.2 Controlled temperature Transcriptome

Our analysis of weather in Ubiaja and Ibadan indicated that day-time temperature Ubiaja was cooler than Ibadan (Figure 1). We hypothesized that the cooler temperatures might be a factor influencing earlier flowering in Ubiaja, and that the genotypic differences in flowering (Figure 5) would relate to their transcriptomes. Under controlled conditions with 22°C as reference, 7253 genes were differentially expressed (5% FDR) in response to the three temperatures studied. 3940 had higher expression and 3313 lower expression at warmer temperatures of 28 and 34°C at a 5% FDR (Figure 8). Notably enriched among genes with higher expression at 28 and/or 34°C were stress responsive processes (p = 1.2 e-17) similar to Ibadan.

**Figure 8.**
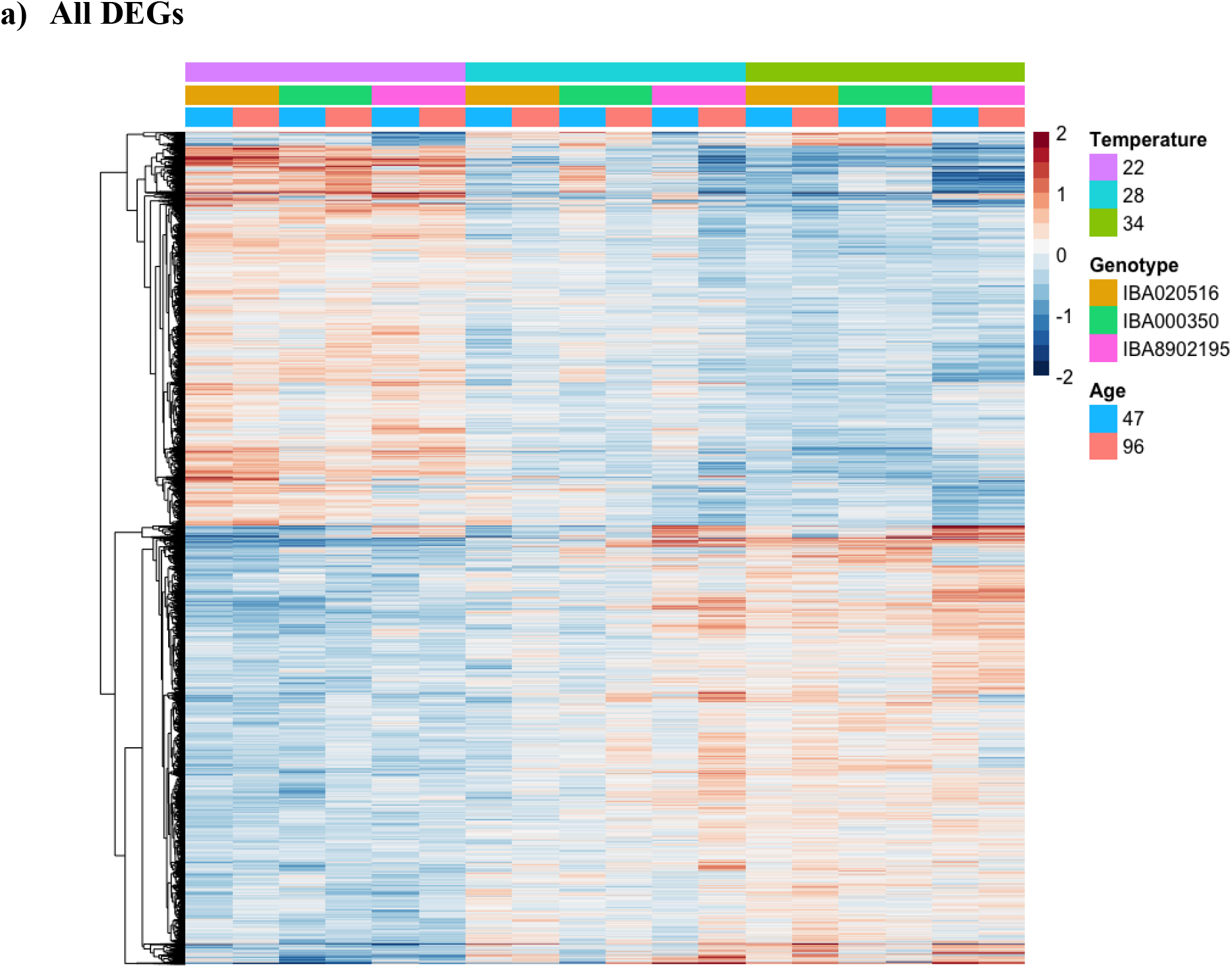
Heat map showing relative expression of differentially expressed genes in temperature (22, 28 and 34°C), genotype (516, 350, 2195), and timepoint of development (47 and 96 dap). Figure shows 7253 genes significantly differentially expressed (α = 0.05) (averages of biological replicates per time point, genotype and location). Colour scale indicates log2 Fold Changes.

##### Flowering time genes

Ninety-six known flowering time genes were differentially expressed under controlled temperature, split nearly evenly between positive and negative effectors, 49 and 47 genes respectively (Figure 9a,b). The flowering enhancing genes (based on characterization in Arabidopsis) GA2ox1, SPL3, LNK1, PRR8, PGM1, FUL, ADG1 and LNK2 had higher expression at 22°C than at warmer temperatures (28°C and 34°C) for both timepoints (47 and 96 dap) (Figure 9a). Most genes known to negatively influence flowering time in Arabidopsis had lower expression at 22°C than at warmer temperatures. Some of these genes had higher expression under 22°C (Figure 9b).

**Figure 9.**
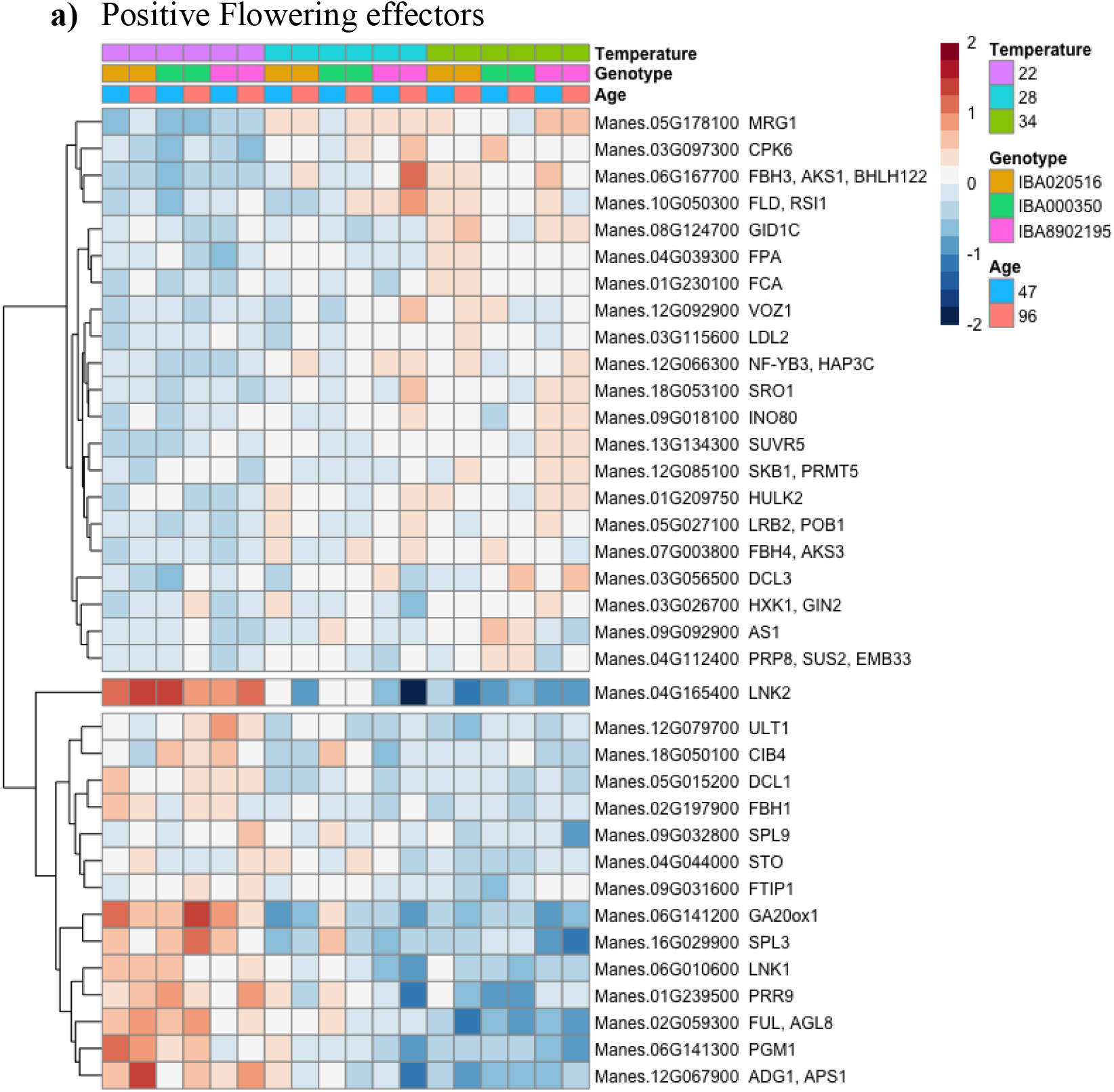
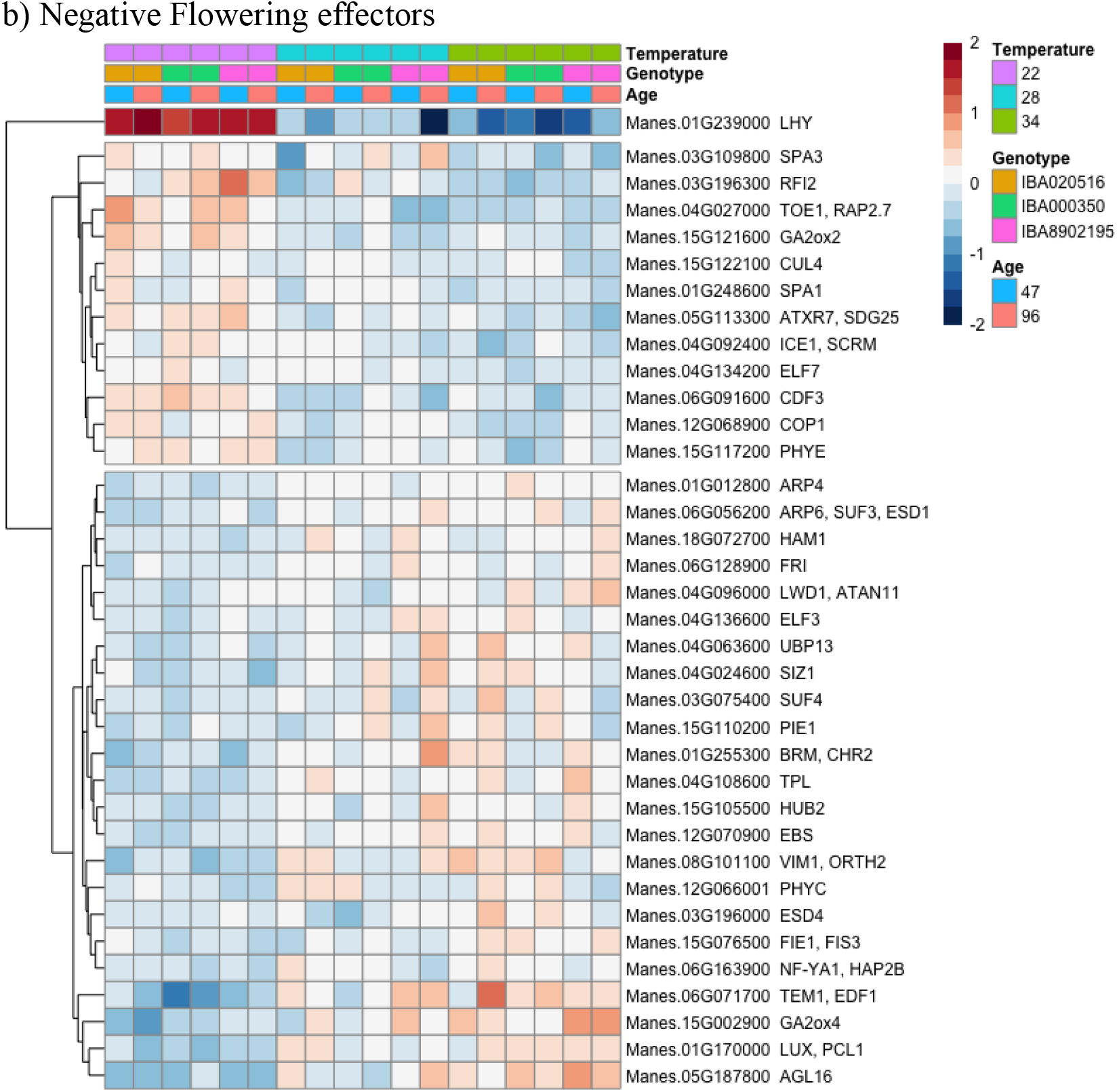
Flowering time genes differentially expressed in the growth chamber study in response to temperature, genotype, and time of development. The heat map shows relative expression across both times of development. a) Positive flowering genes at b) Negative flowering genes

##### Hormone signaling genes

Most hormone signaling genes (from a selected list, see Materials and Methods) differentially expressed under controlled temperature had lower expression at 22°C and higher expression at warmer temperatures (Figure 10). Just like in the field experiment, several abscisic acid (OST1, ABI1, SNRK2-8, AREB3) and auxin (SAUR-like, IAA16, ARF7, IAA30, IAA29, GH3.9) related genes were differentially expressed. In addition, other hormone signaling involve in stress like jasmonic acid signaling genes (JAS1, JAZ12), GA receptor (GID1C), bzip transcription factors involved in multiple hormone signaling pathways (TGA1,PAN), and translation terminator ERF1-3 were differentially expressed. Also, the negative regulator of ethylene stress-hormone pathway, ethylene receptor ETR2, had higher expression at 22°C. Cytokinin signaling was regulated in the direction of suppressed signaling at 22°C: the cytokinin receptor AHK2 and A-type response regulators (ARR8 and ARR9), which function as negative regulators of cytokinin signaling, had higher expression levels at 22°C, whereas B-type cytokinin response regulators (ARR12 and ARR2) mediating cytokinin positive effects had lower expression levels at 22°C.

**Figure 10.**
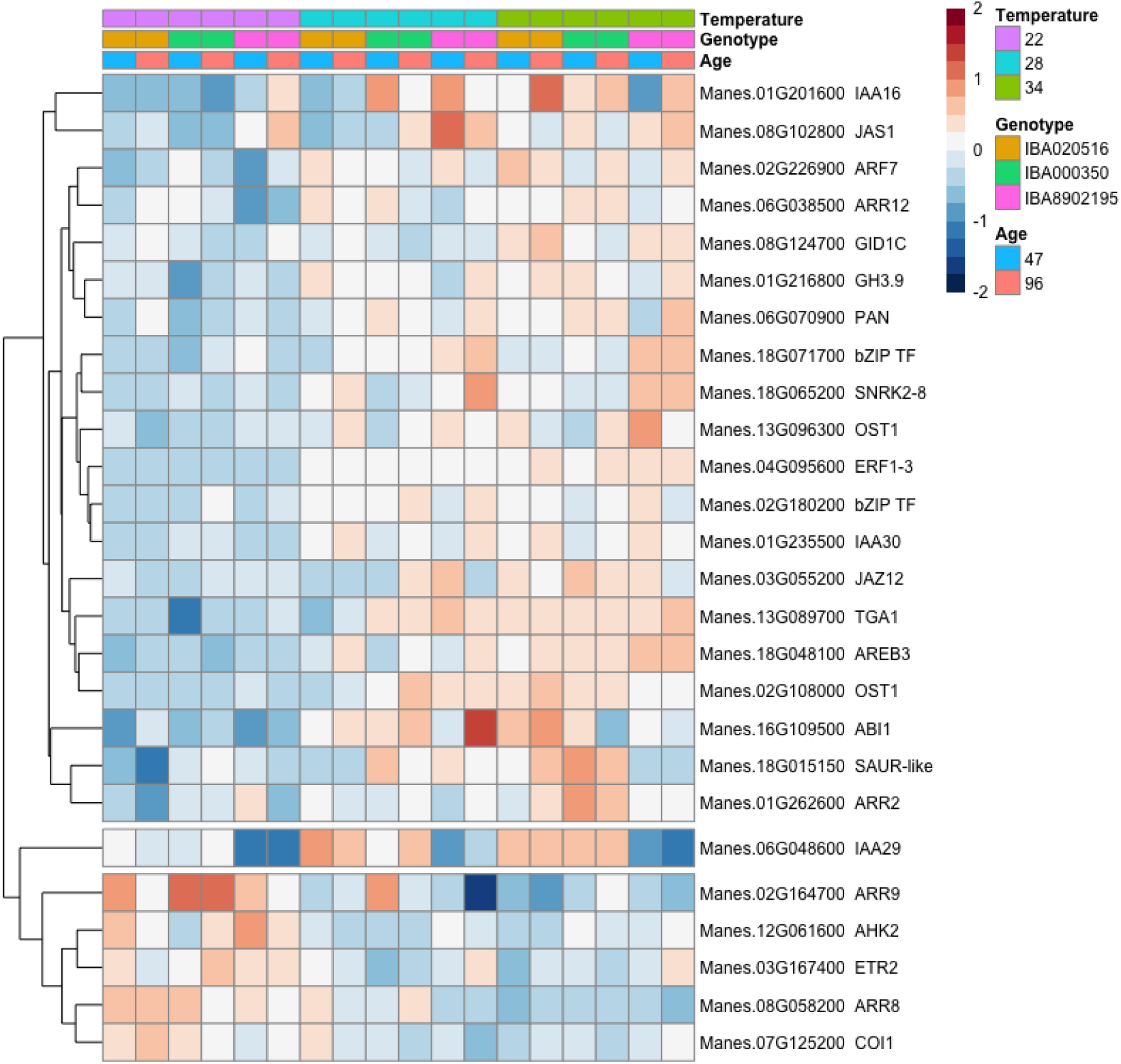
Select hormone signaling genes differentially expressed in the growth chamber study in response to temperature, genotype, and time of development. The heat map shows relative expression across both times of development.

### 3.4 Comparison of overlapping flowering time and hormone signaling DEGs under field and controlled temperature conditions

We compared the expression profiles of flowering and hormone signaling genes that were differentially expressed under both field and controlled temperatures

Four flowering time genes, TEMPRANILLO1 (TEM1), ULTRAPETALA1 (ULT1), PHYTOCHROME E (PHYE) and CONSTITUTIVE PHOTOMORPHOGENIC 1 (COP1), were differentially expressed (P <0.05) under both field and controlled temperature conditions (Figure 11a,b. TEM1, however, had the most consistent expression profile under field and controlled temperature conditions. In the Ubiaja field and at 22°C, TEM1 had the lowest expression levels for all genotypes and irrespective of timepoint. In the Ibadan field, TEM1 expression levels were highest pre-forking and declined post-forking whereas at 28 and 34°C, expression levels were generally high compared with the cooler 22°C.

**Figure 11.**
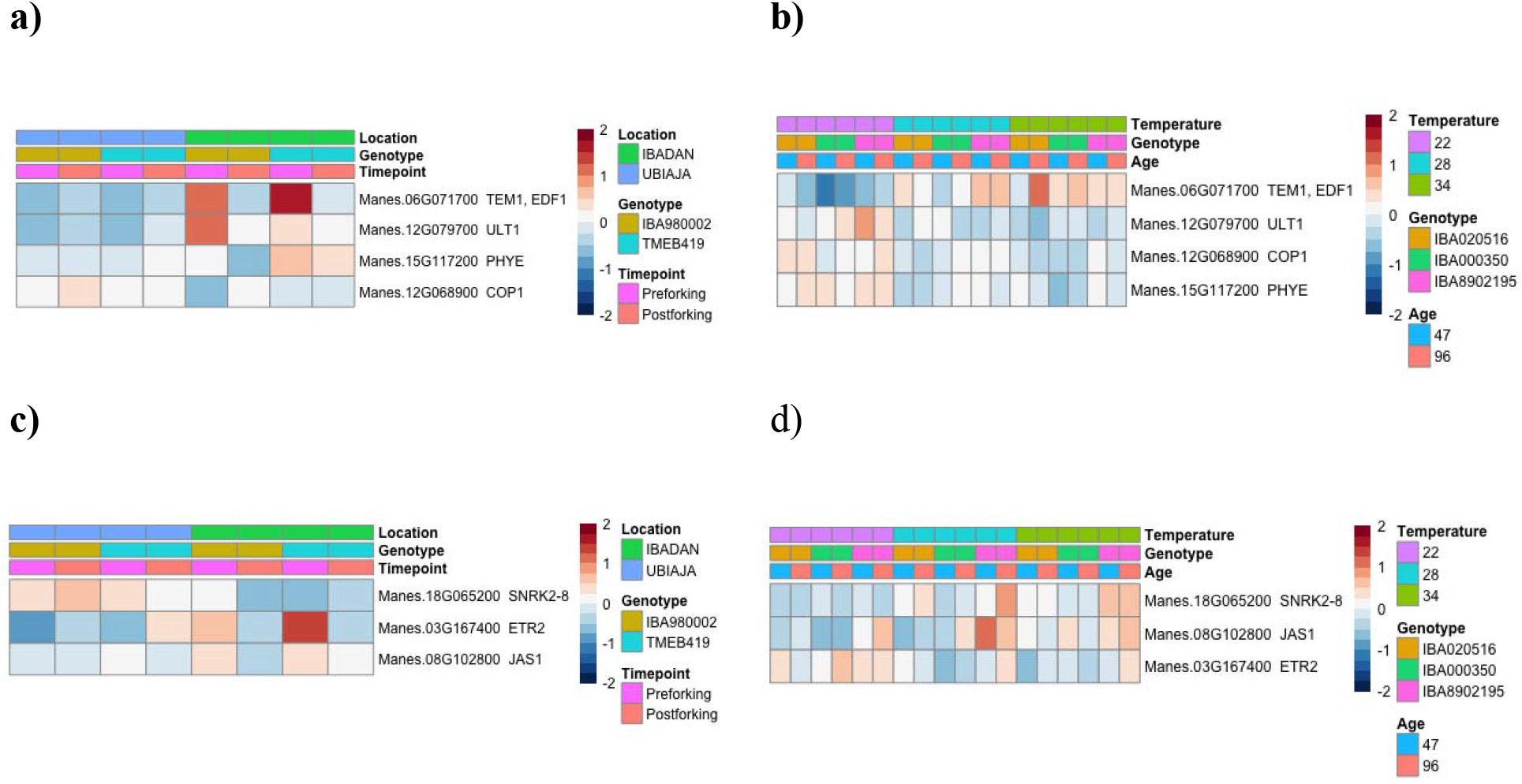
Flowering time and hormone signaling genes commonly expressed between field and controlled temperatures a) flowering time genes on field b) flowering time genes in growth chamber c) hormone signaling genes on field d) hormone signaling genes in Growth chamber.

Three hormone signaling genes, SNRK2-8, ETR2, and JAS1, were differentially expressed under both field and controlled temperature conditions. JAS1 had the most consistent pattern of expression between the field and controlled temperatures. In the Ubiaja field and at 22°C, its expression levels were generally lower that in the Ibadan field and at warmer temperatures (Figure 11c,d). The expression patterns for SNRK2-8 and ETR2 were not consistent between field and controlled temperature given observed flowering time. SNRK2-8 generally had higher expression in Ubiaja, than in Ibadan whereas under controlled temperatures, expression levels were generally lowest at 22°C. ETR2 on the other hand generally had lower expression in the Ubiaja than Ibadan while at controlled temperatures its expression levels were generally higher at 22°C than at warmer temperatures (Figure 11c,d).

## 4 Discussion

### Vegetative growth showed complexity between field and controlled temperatures

The current study and previously published work have indicated that flowering is earlier and flower production is better in Ubiaja than Ibadan [17, 18, 33, 34]. One goal of the present study was to provide insight on the underlying basis of this difference. Plant growth was generally more vigorous in Ibadan than Ubiaja (as evidenced by plant height, shoot weight and storage root numbers). However the partitioning index, showing shoot weight per plant total weight was higher in Ubiaja, suggesting more resource focused on storage root development (Figure 2). Under controlled temperatures, plant vegetative growth response was not linear as observed on the field. Plant height increased with increase in temperature, storage root numbers tended to be higher at 14ooler temperatures and partitioning index tended to be highest at the intermediate temperature (28°C) (Figure 4). Although the temperatures in Ibadan were somewhat warmer than in Ubiaja (Figure 1), the comparison of field and controlled temperature plant growth indicate that temperature alone does not explain the differences between vegetative growth in the Ibadan and Ubiaja field environments. An earlier study by [18] indicated that the percent nitrogen content in Ubiaja soil (0.131) was lower than that in Ibadan soil (0.167). It is known that nitrogen limitation induces plants to invest in root development at the expense of shoots [35] and this is in line with a higher partioning index in Ubiaja (storage root weight/whole plant weight). A possible hypothesis for the earlier flowering in Ubiaja is that the environment in Ubiaja might suppress vegetative growth and as a consequence provide better photosynthate supply flower development. Shoot growth was indeed smallest in Ubiaja and at 22°C (temperature with earliest flowering). The complex relationship between root growth and temperature however challenges this hypothesis. Water status (Rainfall) and differing soil nitrogen levels under field conditions were interacting factors not accounted in controlled temperature study. [36] showed that in cassava root partitioning index was not significantly affected by water limitation; unless water limitation was prolonged to the point of remobilization from stem and root storage reserves [37]. This was contrary to the root partitioning observed in Ubiaja as it received more rainfall. Previously [18] suggested that the difference in soil type or fertility did not explain the considerably better flowering in Ubiaja. The effect of the interaction between temperature, water status and soil nutrient on cassava flowering time should be investigated.

### Early flowering genotypes were relatively insensitive to the environment while in late flowering genotypes, the Ibadan environment and warmer temperatures had a delaying effect

Genotypes such as ‘0002, ‘275, ‘615 and ‘516 flowered early and had fewer nodes to forking (about 18 or less under field conditions or about 30 or less under controlled temperatures). The number of nodes to forking was independent of plant height (Figures 2 - 5). The flowering rates were similar for these genotypes in Ubiaja and Ibadan fields with a 0.5 probability corresponding to a chronological age of about 60 to 70 dap. Similarly, under controlled environments with day-time temperatures between 22 and 34°C, the 0.5 flowering probability was also about 70 dap in ‘516.

Genotypes such as ‘350, ‘2195, ‘085 and ‘419 showed very large responses to the environment with respect to their flowering time and were especially late in the Ibadan field and at warmer temperatures. They had significantly more nodes to forking in Ibadan and higher probability of not flowering when grown at warmer temperatures (28°C and 34°C), compared to the Ubiaja field and at the cooler 22°C. In Ubiaja or at 22°C, the number of nodes to flowering in late-flowering lines were reduced to values approaching those of early flowering genotypes under those conditions (Figures 3, 5).

A meta-analysis of flowering time data on of over 700 genotypes in grown at Ubiaja and Ibadan [33] showed that modal flowering time in both locations was also between 60 and 70 dap (Figure S2). It is therefore likely that the flowering times of early genotypes represent the minimal or most probable flowering time of cassava in the absence of environmental conditions such as warmer temperatures that induce regulatory systems which delay flowering. Our studies indicate that later genotypes primarily differ from early ones in the extent to which their flowering is delayed in unfavorable environments, i.e. Ibadan and warm growth chambers.

The overexpression of Arabidopsis FT in cassava [4, 16] and of a native FT gene in cassava [15] resulted in significantly earlier flowering in late cassava genotypes. These early flowering phenotypes were accompanied by significantly reduced number of nodes to forking [4, 15], thus confirming that earliness is associated with a reduction number of nodes to fork type branching. Furthermore, [5] showed that the late-flowering genotype ‘419 initiates flowers at 22°C but not at warmer temperatures, which is in agreement with the current findings.

Several members of the Euphorbiaceae family, to which cassava belongs, are known to flower more readily at moderately cool temperatures than at warmer temperatures, including rubber tree (Hevea brasiliensis) [38] poinsettia (Euphorbia pulcherrima) [39], and leafy spurge (Euphorbia esula) [40]. Other tropical perennials are also known to be induced to flower by cool ambient temperature, notably Lychee (Litchi chinensis) and Mango (Mangifera indica). In Lychee, warm temperatures stimulate vegetative growth while cool temperatures of 20°C or less promote reproductive growth [41, 42]. In mango, cool temperatures of 15 °C stimulated flowering [43]. Furthermore, in mango, water stress at cool temperatures causes profuse flowering but water stress under warm temperatures did not induce flowers [43]. This stimulation of flower induction by cool temperature in the tropics has been suggested to be related to the drop in temperature preceding the onset of rains, thus serving as an environmental cue [44].

### Flowering repressors are highly expressed in Ibadan before forking

The current study determined the transcriptome of expressed genes in recently matured leaves of the Ibadan-Ubiaja field experiment, and of the temperature comparison in the growth chamber experiment. In the favorable-flowering Ubiaja environment, cassava homologues of known Arabidopsis flowering repressors, including GIBBERELLIC ACID 2 OXIDASE 1 (GA2ox1), GIBBERELLIC ACID 2 OXIDASE 8 (GA2ox8), TEMPRANILLO 1 (TEM1) and PHYTOCHROME E (PHYE) [13] generally had low expression levels before and after forking. In contrast, these genes were highly expressed in poor-flowering Ibadan environment before forking, but their expression declined after forking. On the other hand, a cassava homolog Flowering Locus T (MeFT1), was generally expressed at higher levels before in Ubiaja than Ibadan. [5] first established that MeFT1 expression was related to flowering tendency as it was expressed at higher levels in ‘0002 (early genotype) than in ‘419 (late genotype) while the over expression studies of MeFT1 by [15] further confirmed its florigenic properties. The role of FT as a mobile long distance signaling peptide that moves from leaf to shoot apical meristem to induce flowering has long been established in Arabidopsis [14, 45]. The current expression profiles are sensible for the earlier flowering times in Ubiaja relative to Ibadan (Figure 3) as it corresponds with the continuously low expression of flowering time repressors and relatively higher expression of a florigen at both developmental stages studied.

In Ibadan, the expression profiles of flowering repressor genes correlated well with developmental stage but the florigen generally had low expression at all timepoints. In this case flowering may be promoted by obtaining an optimal ratio between flowering enhancers and repressors rather than only an increased expression of enhancers. This observation is in line with a previously described model in tomatoes, in which flowering, and plant architecture is determined by the local balance of florigenic and anti-florigenic signals in respective organs [46, 47].

### Expression of some cassava homologues of flowering time genes correlates with temperature rather than flowering response

Under both field and controlled temperature conditions, some floral regulatory genes were expressed in a direction contrary to what was expected for their role in flowering (Figure 7, Figure 9). Some flowering repressors had higher expression levels in Ubiaja and at 22°C, which are conditions at which cassava flowering is earlier e.g., CONSTITUTIVE PHOTOMORPHOGENIC 1 (COP1). Some flowering enhancers had higher expression levels in Ibadan and at warmer temperatures which had a delaying effect on cassava flowering time. It is notable that a majority of the known flowering time genes were assigned a positive or negative role based on the flowering time phenotype of Arabidopsis mutants. While both Arabidopsis and cassava are long day plants [5, 12], Arabidopsis flowers earlier at warm ambient temperatures [48] while cassava flowers earlier at cooler temperatures [5].

As an example, both LHY and COP1 have been defined as flowering repressors in Arabidopsis. [49, 50]. But these genes are known to be involved in plant temperature sensing and thermomorphogenesis [51, 52]. The distinct temperature response profiles of these genes which did not correlate with flowering response may reflect their activity cassava’s perception of the environment.

### Expression profiles of hormone signaling genes responds to plant growth environments

Several Abscisic Acid signaling genes were modulated in response to field environments and controlled temperatures, genotypes and developmental stages. The SNRK2-8 in particular was differentially expressed under both field and controlled temperatures but the pattern of expression was however very complex. SNRK2-8 had higher expression levels in Ubiaja, compared to Ibadan and lower expression levels at 22°C compared to warmer temperatures 28 and 34°C (Figure 11). SNRK2-8 phosphorylates and thus activates other ABA response genes [53, 54]. Apart from Abscisic acid signaling, other hormone genes were modulated but notable is the Ethylene receptor ETR1 and Jasmonic signalling JAS1. Like SNRK2-8, ETR1 showed a complex expression pattern between field and controlled temperatures. The expression pattern of JAS1 was the most consistent between field and controlled temperatures. These complex expression profiles possibly reflect plant adaptation to growth environment as needed.

### Flowering phenotype correlates with TEM1 expression under field and controlled temperature conditions

Flowering time genes, TEM1 (the Arabidopsis homologue specifically on chromosome 6 in cassava), and COP1 had the most similar expression patterns between field and controlled temperatures environments (Figure 11). The TEM1 expression pattern, however, was most correlated with observed flowering times under all environmental conditions. Flowering repressor TEM1 had low expression levels under all conditions in which cassava flowering is earlier for all genotypes (i.e. Ubiaja environment and 22°C) (Figure 3 and 5). This low expression was observed early in plant life (before forking in the field and at 47d in the growth chamber) and was maintained even after forking on the field or at 96d in the growth chamber (when at least 70% of all genotypes had forked). In Arabidopsis, TEM1 has a role in regulating juvenility [55]. So low expression levels early in plant life will reduce length of juvenile phase as seen in our study. TEM1 also directly represses FT expression under conditions that delay flowering – in TEM1 knock out mutants, FT expression was consistently higher than wild type while in overexpression lines there was barely any FT expression [56]. In the field, MeFT1 expression was generally higher when the expression of TEM1 was low in line with [56]. In the growth chamber both FT homologues in cassava were not significantly differentially expressed in line with [5] observation that cassava FTs were not be clearly temperature responsive. The expression pattern of indicates that it is an important flowering inhibitor in cassava. Its relationship with FT in cassava especially under controlled temperatures merits further investigation.

## 5 Conclusion

We have analyzed the flowering time and transcriptome of cassava under field and controlled conditions and found that in the Ubiaja field conditions and cool ambient temperatures of about 22°C cassava flowered early. Late flowering genotypes were much more sensitive to their growth environments than early flowering genotypes and their delayed flowering time was pronounced in the Ibadan field and at warmer temperatures. The transcriptomes we revealed under field and controlled-temperature conditions indicated that some flowering time genes were expressed in a temperature dependent manner rather than in relation to a flowering time. The flowering repressor gene TEM1 had consistently low expression levels under conditions in which cassava flowering time was early (i.e. at 22°C and at Ubiaja) indicating that it is an important flowering inhibitor in cassava.

