## Supplementary figures for "Environmental responsiveness of flowering time in cassava genotypes and associated transcriptome changes"

a)

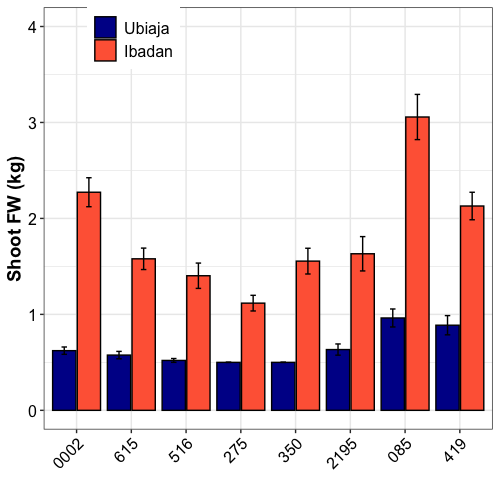

b)

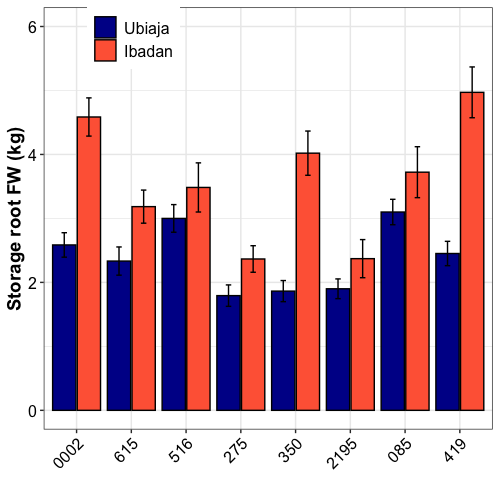

Figure S1. Shoot and storage root Fresh Weight under field conditions (a) Shoot Fresh Weight (kg) (b) Storage root Fresh Weight (kg)

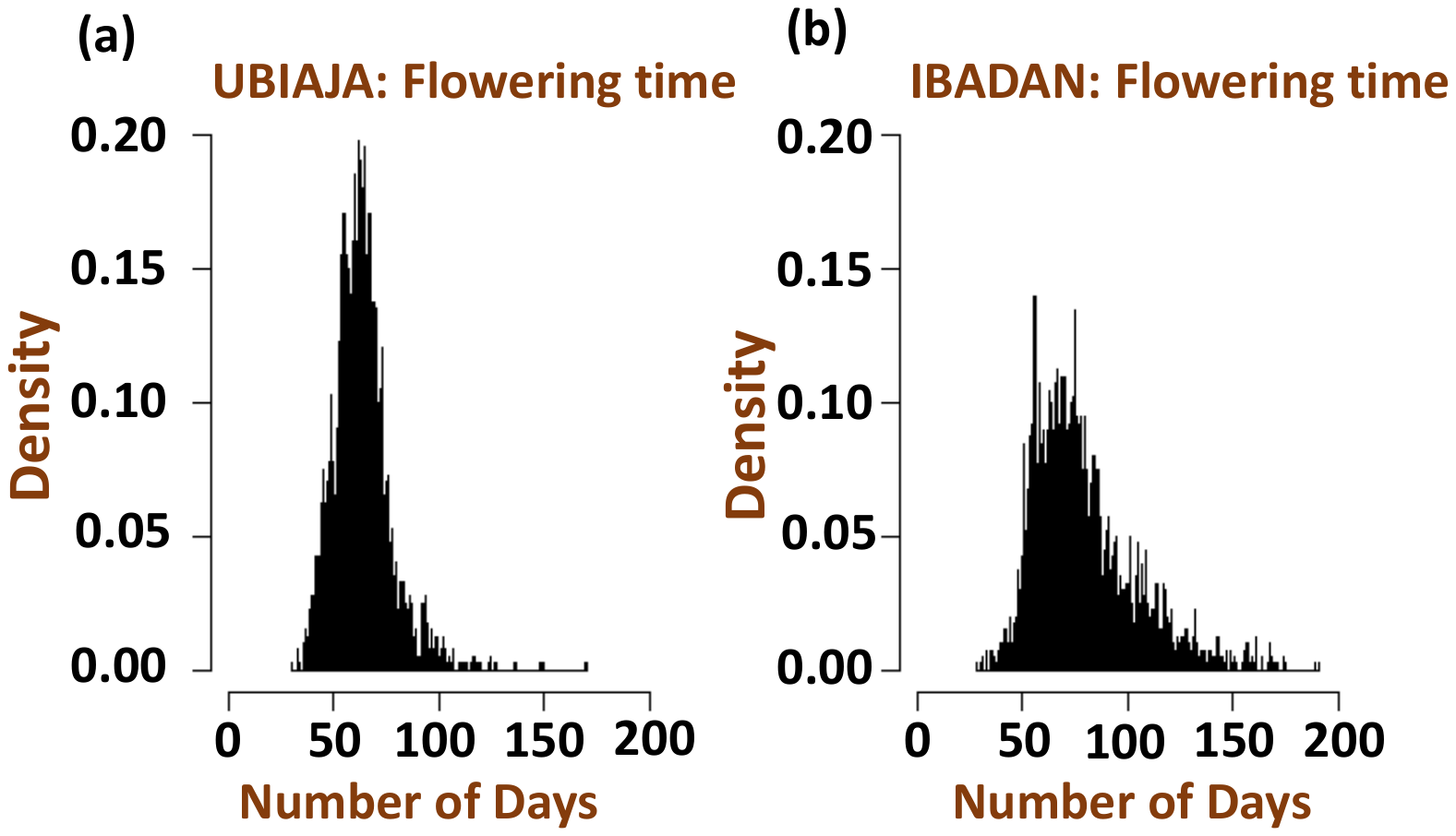

**Figure S2** Meta analysis of the distribution of flowering times in IITA’s diversity population (Genetic Gain) between 2013 and 2016. Source – Diebiru (2017). a) Ubiaja b) Ibadan

a

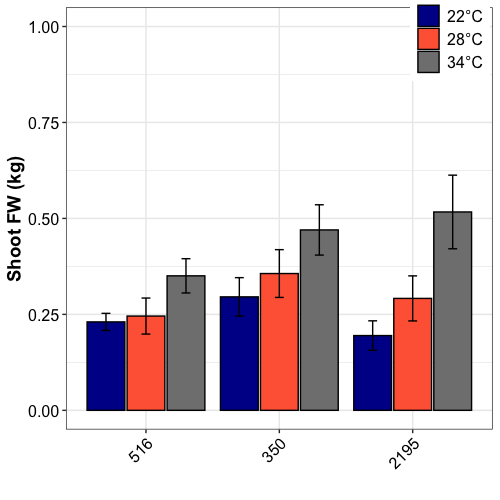

b

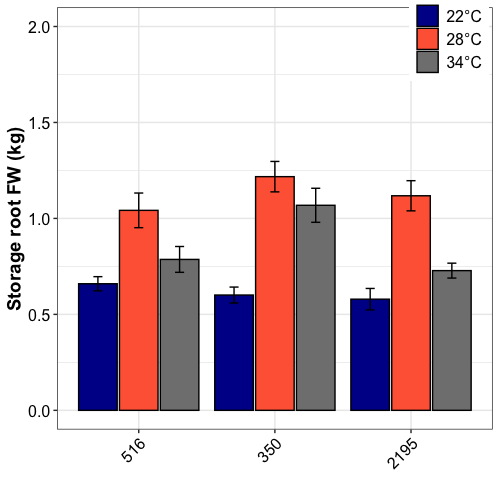

Figure S2. Shoot and storage root Fresh Weight under controlled temperature conditions (a) Shoot Fresh Weight (kg) (b) Storage root Fresh Weight (kg)
